# Identification and Credentialing of Patient Derived Xenograft Models of Invasive Lobular Breast Carcinoma using Multi-omics and Histopathology assessment

**DOI:** 10.1101/2025.08.21.669685

**Authors:** Jagmohan Hooda, Jennifer M Atkinson, Osama Shiraz Shah, Megan Yates, Daniel D Brown, Morgan DeBerry, Stefano Cairo, Paolo Schiavini, Hsiu-Wen Tsai, Marianna Zipeto, Rohit Bhargava, Steffi Oesterreich, Adrian V Lee

**Affiliations:** Women’s Cancer Research Center, University of Pittsburgh Medical Center (UPMC) Hillman Cancer Center (HCC), Magee-Womens Research Institute, Pittsburgh, PA; Department of Pharmacology & Chemical Biology, University of Pittsburgh, Pittsburgh, PA; Institute for Precision Medicine, University of Pittsburgh, Pittsburgh, PA; Department of Pathology, University of Pittsburgh, Pittsburgh, PA; Champions Oncology, Hackensack, NJ

**Keywords:** Invasive Lobular Breast Cancer, Patient-Derived Xenograft, *CDH1*, Genomic Profiling, Targeted Therapy

## Abstract

Breast cancer is a heterogeneous disease with numerous histological subtypes. Invasive lobular cancer (ILC) is the most common special subtype, accounting for 10-15% of all breast cancers. The pathognomonic feature of ILC is the loss of E-cadherin (CDH1), which leads to a unique single-file growth pattern of discohesive cells. Although ILCs show better prognostic factors than the most common No Special Type (NST) of breast cancer, patients with ILC have worse long-term outcomes, which is not well understood. In this study, we aimed to identify and characterize Patient-Derived Xenograft (PDX) models of ILC based upon the presence of truncating *CDH1* mutations and/or low *CDH1* mRNA expression among 128 human breast cancer PDX models. We selected 8 PDX models for validation using Immunohistochemical (IHC) analysis for E-Cadherin, p120, ER, PR, and HER2. We confirmed that seven of these PDX models are indeed ILC while one was identified as mixed NST-ILC PDX. Molecular analysis of the confirmed ILC PDX models showed enrichment of truncating *CDH1* mutations, significantly lower levels of *CDH1* mRNA expression and predominantly luminal subtypes compared to NST PDX models, in line with the molecular characteristics of human ILC disease. The commonly altered genes in the ILC PDX models included *PIK3CA* (57%), *CDH1* (57%) and *TP53* (57%) among others. Our study confirms and characterizes new ILC PDX models, offering valuable tools to advance our understanding of human ILC biology and support the development of innovative treatment strategies.

## Introduction

Breast cancer is a heterogeneous disease with numerous histological subtypes. The most common subtype is No Special Type (NST), accounting for more than two-thirds of all breast cancers. Invasive Lobular Cancer (ILC) is the most common special subtype, accounting for 10-15% of all breast cancers [1, 2]. In the US alone, there will be an estimated 47,500 new cases of ILC in 2025, which represents approximately 15% of all invasive breast cancer diagnosis [1, 2]. The pathognomonic feature of ILC is the loss of E-cadherin (encoded by the *CDH1* gene) and cytoplasmic p120, which leads to a lack of adherens junctions and a unique single-file growth pattern of discohesive ILC cells [3, 4]. This growth pattern decreases detection by mammography, resulting in late detection and larger tumors. Although ILCs show similar or even better prognostic factors than NST, patients with ILC have worse long-term outcomes [1, 2, 4-6].

ILC has historically been understudied, in part due to the lack of appropriate research models [6, 7]. Since ILC affects “only” 10-15% of all breast cancers, the majority of models have been generated from the more frequent subtype, NST or invasive ductal carcinoma (IDC) [4]. For example, the Cancer Cell Line Encyclopedia (CCLE) contains 54 NST cell lines but only 2 ILC cell lines. Additionally, only a limited number of patient-derived xenograft (PDX) models are evident in the published literature. There is a critical need for additional in vitro and in vivo models to study ILC biology, as well as to test targeted therapies. ILC PDX and patient-derived xenograft organoids (PDXO) are particularly valuable tools to enable target validation and assess drug treatment response [8].

The purpose of this study was to identify and characterize PDX models of ILC. Specifically, we used Whole-Exome Sequencing (WES) and RNAseq data from 128 human breast cancer PDX models to identify putative ILC PDX models based upon the presence of truncating *CDH1* mutations and/or low *CDH1* mRNA expression. Next, we validated a subset of these potential models by Immunohistochemistry (IHC) analysis for E-Cadherin, p120, ER, PR, and HER2 and confirmed seven of these PDX models as ILC. The findings from this study provide evidence for new PDX models that may be utilized to provide insights into the molecular characteristics of human-ILC disease and develop new treatment strategies.

## Methods

### Patient derived xenograft models

Champions Oncology is a preclinical research and clinical specialty testing provider that has developed over 1,400 human PDX models [9, 10]. Champions Oncology obtains consented patient samples to engraft, develop and characterize PDX models from a range of tumor types, primarily in immunocompromised mice. We queried the Champions’ collection of breast cancer models (n=128) for potential PDX models of ILC.

### Bioinformatics analysis

To identify and confirm new ILC PDX models, we downloaded the mutation, copy number and RNA expression data on all breast cancer PDX models (N=128) in the Champions Oncology’s LUMIN [9, 10] portal on February 22^nd^, 2023. All downstream analysis were performed using R. PAM50 molecular subtypes were computed from RNA expression data using molecular subtyping function from R package genefu [11]. Mutation calls were filtered to exclude silent mutations. Copy number calls were defined as GAIN (copy number = 1), LOH (copy number = −1), DIPLOID (copy number = 0), AMP/AMPLIFICATION (copy number = 2) and DEL/DELETION (copy number = −2). Putative ILC models were defined based on the following criteria: truncating *CDH1* mutations and/or low *CDH1* mRNA. We also included models with a clinical annotation of lobular disease of the corresponding clinical samples. Based on availability of PDX tissue and resources, we selected N=8 potential ILC PDX samples for further validation using immunohistochemical and histomorphic analysis. For further details regarding the genomic data, please contact Champions.

### Immunohistochemistry

FFPE tissue from selected breast cancer PDX models was sectioned and subjected to immunohistochemical (IHC) analysis of estrogen receptor (ER), progesterone receptor (PR), HER2, E-Cadherin and p120. Selected models were assessed at the earliest passage/transplant generation with FFPE material available for analysis as follows: CTG-1714 (P3+1), CTG-2432 (P2+1), CTG-2518 (P2+1), CTG-2611(P3+1), CTG-2810(P4+1), CTG-2849 (P3), CTG-2930(P5+1), CTG-3283 (P4), CTG-3399 (P4), CTG-3434 (P3 and P5). The antibodies and the protocol used in this study were as follows: E-cadherin (Clone: 36; Vendor: Ventana, Tucson, AZ; Dilution: ready to use [RTU], Pre-treatment: CC1-S, Detection: Ultraview; Staining platform: Ventana Benchmark Ultra), p120 (Clone: 98; Vendor: BD Biosciences, Franklin Lakes, NJ; Dilution: 1:200; Pre-treatment: CC1-S; Detection: Ultraview; Staining platform: Ventana Benchmark Ultra), ER (Clone: SP1; Vendor: Ventana, Tucson, AZ; Dilution: RTU, Pre-treatment: CC1-S, Detection: Ultraview; Staining platform: Ventana Benchmark Ultra), PR (Clone: 1E2; Vendor: Ventana, Tucson, AZ; Dilution: RTU, Pre-treatment: CC1-S, Detection: Ultraview; Staining platform: Ventana Benchmark Ultra), and HER2 (Clone: 4B5; Vendor: Ventana, Tucson, AZ; Dilution: RTU, Pre-treatment: CC1-M, Detection: Ultraview; Staining platform: Ventana Benchmark Ultra). Stained slides were interpreted by a trained breast pathologist (RB).

## Results

We acquired WES and RNAseq on 128 breast cancer PDX models from the Champions Oncology’s LUMIN portal. Principal component analysis of the RNAseq dataset, using the top 10% variably expressed genes, revealed two major clusters, as expected, representing the Basal and Luminal intrinsic molecular subtypes (***Supplementary Figure S1A***). We then identified potential ILC PDX models by examining their E-cadherin mRNA expression and *CDH1* gene mutations. Using RNAseq data, we identified 11 putative ILC PDX models with low E-cadherin mRNA levels (z-score <= −0.5) (***Figure 1A***). Separately, using WES data, we identified 9 ILC PDX models with various types of non-recurring *CDH1* mutations (truncating, missense, and others) distributed across the gene body (***Figure 1B***). Among these, there was an overlap of 7 PDX models that had both low *CDH1* levels and non-recurring *CDH1* mutations (***Supplementary Table S1***). As expected, models with truncating *CDH1* mutations exhibited the lowest *CDH1* mRNA expression (***Figure 1C***), a characteristic enriched in human ILC [12, 13]. We flagged 4 PDX models with truncating *CDH1* mutations as putative ILC PDX models, of which 2 have low *CDH1* levels and two have mid-levels of *CDH1* (***Supplementary Table S1***). In total, we identified 13 potential ILC PDX models using a combination of criteria including truncating *CDH1* mutations and/or low *CDH1* mRNA expression (***Supplementary Table S1)***. Based on tissue availability and resources, we selected 8 putative ILC PDX models for further histological and immunohistochemical (IHC) analysis (***Supplementary Table S2***). Notably, 3 of these putative ILC PDX models were annotated as being generated from human ILC, while the remaining 5 were classified as NST (i.e., ‘carcinoma’ or ‘ductal carcinoma’). Additionally, we selected 2 randomly chosen NST PDX models for comparison.

**Figure 1:**
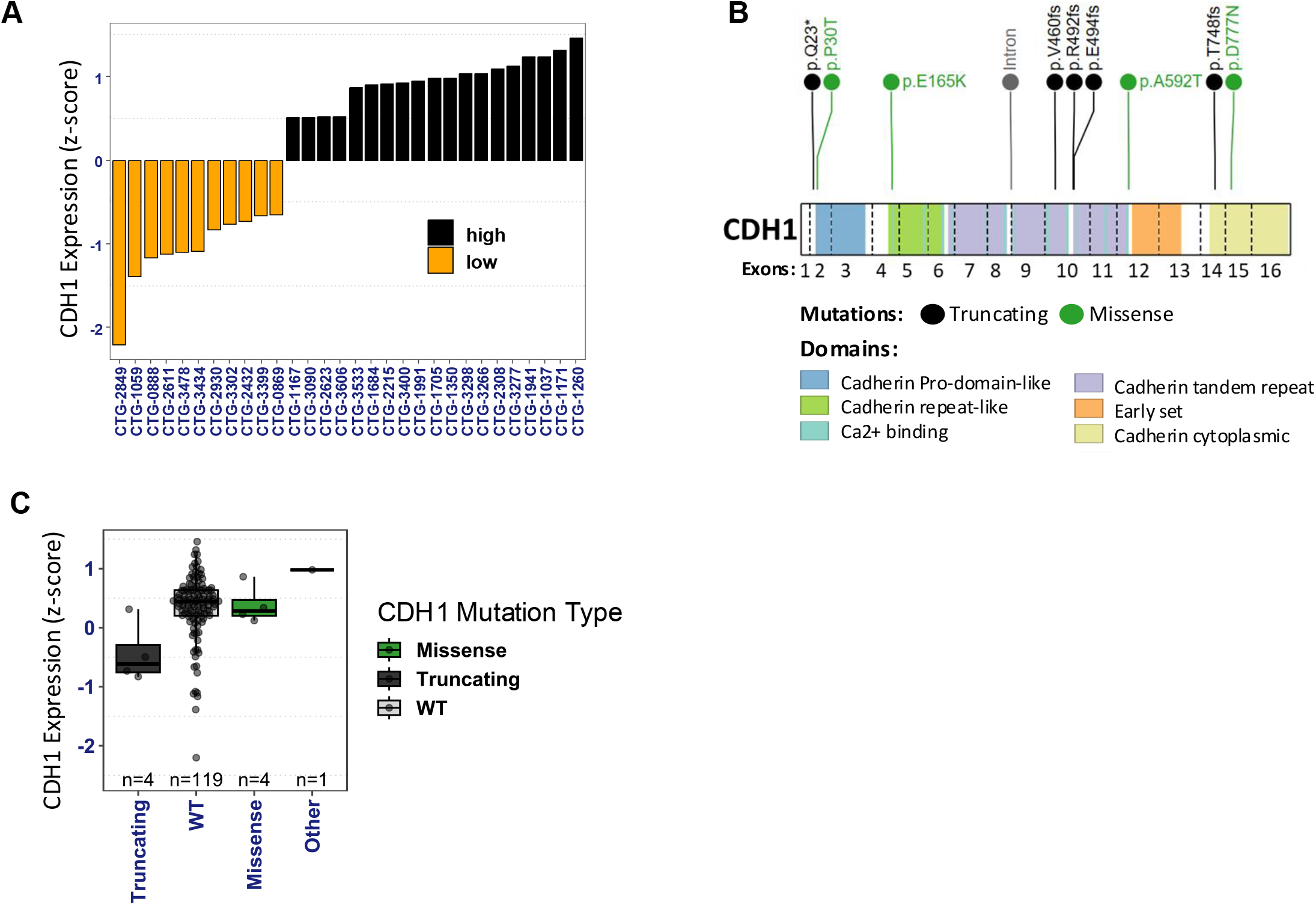
Identification of potential ILC PDX models using multiomic analysis of Breast Cancer PDX models. **A)** Bar plot showing PDX models with highest and lowest *CDH1* mRNA level (z-score) categorized discretely as high (z-score > 0.5), medium/mid (−0.5 < z-score <= 0.5) and low (z-score <= −0.5). The low level PDX models, many of which were classified as ‘carcinoma’, were flagged as putative ILC models. **B)** Lollipop plot showing mutations across *CDH1* gene body. **C)** *CDH1* expression in PDX models with *CDH1* mutations of various types (truncating, missense and other (intronic)) and or wild type (WT) status. All PDX models with truncating *CDH1* mutations were flagged as putative ILC models.

The ten PDX models were subjected to immunohistochemistry (IHC) analysis and pathology review for E-Cadherin, p120, ER, PR, and HER2 (***see Table 1 and Figures 2***). IHC results showed that five of the candidate ILC models (CTG-2432, CTG-2849, CTG-2930, CTG-3283, and CTG-3399) exhibited a loss of E-cadherin expression and cytoplasmic localization of p120. Two further models (CTG-2611 and CTG-2810) showed aberrant membranous E-cadherin, but this was associated with cytoplasmic p120, indicating dysfunctional adherens junction. The remaining putative ILC model, CTG-3434, had an incomplete pattern of E-cadherin loss consistent with mixed NST ILC feature. Because of the mixed histology of CTG-3434 we attempted to grow this model in mice however the PDX failed to grow. In addition, Champions Oncology reported that this generation (P3) has a lower take rate however, a later generation P5 grew successfully. Therefore, we analyzed tissue from the P5 generation and upon IHC analysis, P5 generation was also identified as mixed NST ILC (***Figure 2B***). As expected, the two NST control models (CTG-1714 and CTG-2518), exhibited membranous E-cadherin and p120 (***Figure 2C***).

**Table 1:** Summary of histopathology analysis of the selected PDXs.

| PDX Model # | Tumor markers<br>(PDX) | Selection<br>Criteria | E-cadherin IHC<br>(PDX) | P120 IHC<br>(PDX) | Histopathology<br>Evaluation |
| --- | --- | --- | --- | --- | --- |
| CTG-2432 | ER+/PR+/HER2 2+<br>(HER2 non-amplified) | Truncating CDH1<br>Mutation | Negative | Cytoplasmic | Lobular |
| CTG-2611 | ER+/PR+/HER2 1+ | Low CDH1<br>mRNA | Negative/aberrant<br>(lobular pattern) | Cytoplasmic | Lobular |
| CTG-2810 | ER+/PR-/HER2 1+ | Truncating CDH1<br>Mutation | Weak membranous<br>(lobular pattern) | Cytoplasmic | Lobular |
| CTG-2849 | TNBC | Low CDH1<br>mRNA | Negative | Cytoplasmic | Lobular |
| CTG-2930 | ER+/PR+/HER2 1+ | Truncating CDH1<br>Mutation | Negative | Cytoplasmic | Lobular |
| CTG-3283 | ER+/PR-/HER2 2+<br>(HER2 non-amplified) | Truncating CDH1<br>Mutation | Negative | Cytoplasmic | Lobular |
| CTG-3399 | TNBC | Low CDH1 mRNA | Negative | Cytoplasmic | Lobular |
| CTG-1714 | TNBC | *NST comparison* | Membranous | Membranous | Ductal |
| CTG-2518 | TNBC | *NST comparison* | Membranous | Membranous | Ductal |
| CTG-3434 | ER+/PR- (<1%)/HER2 1+ | Low CDH1<br>mRNA | 70% negative, 30%<br>membranous | 70% cytoplasmic,<br>30% membranous | Mixed NST ILC |

**Figure 2:**
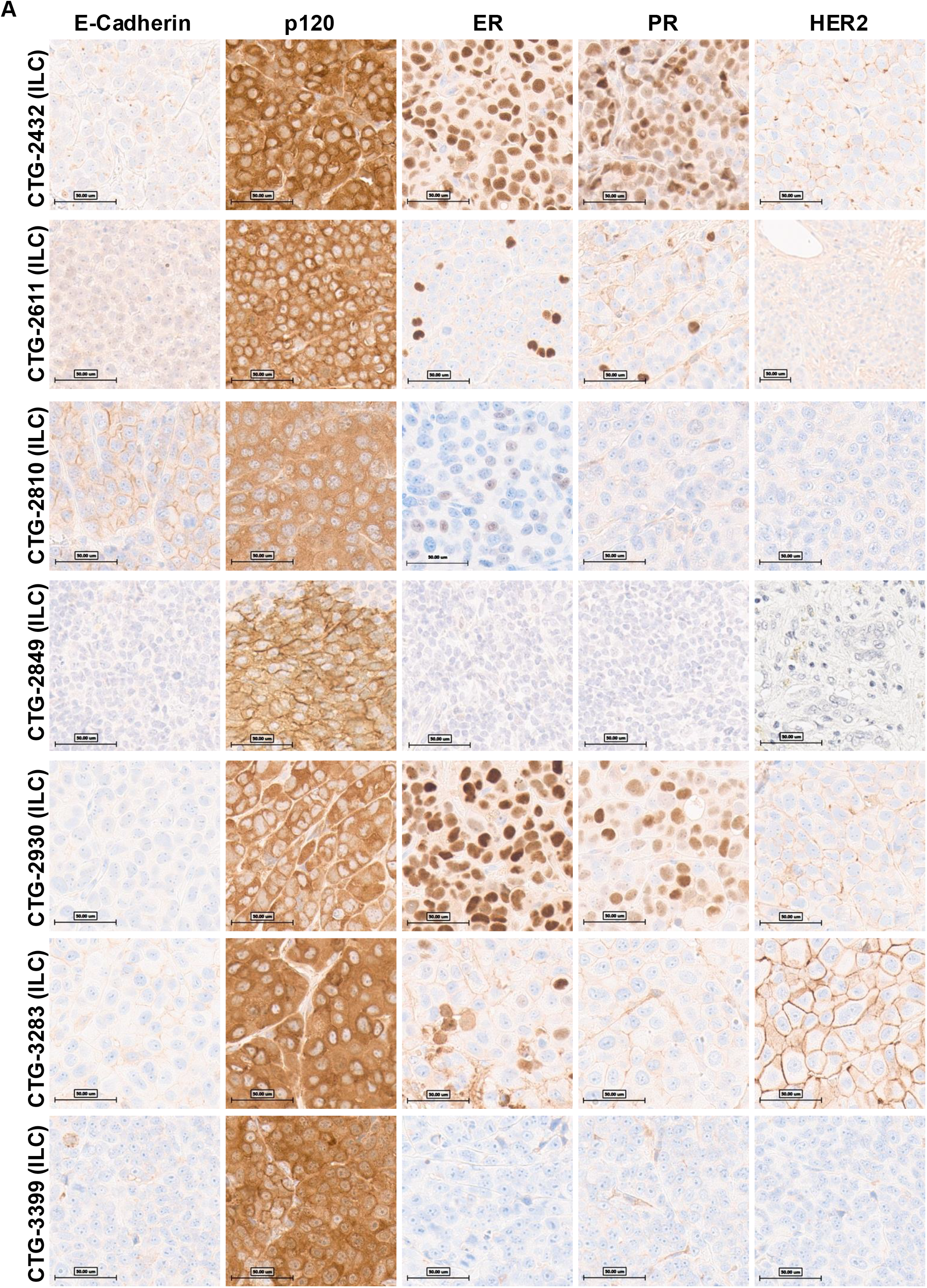
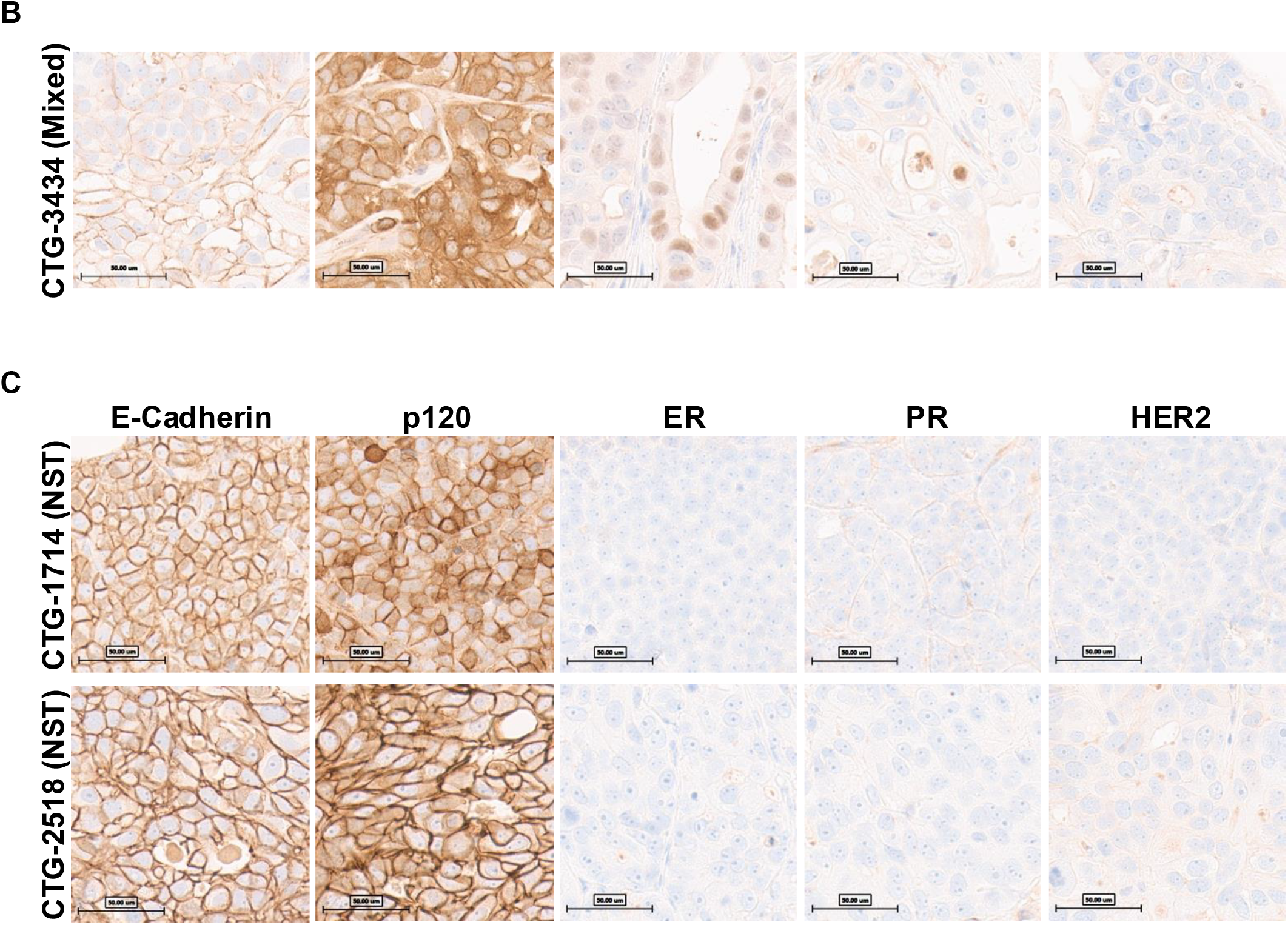
Pathology assessment of PDX models identifying 7 ILC PDX models in Champions database. The PDX FFPE slides underwent staining with E-Cadh, p120, ER, PR, or Her2 antibodies on the Ventana Benchmark Ultra staining platform, and signal detection was conducted using Ultraview. A trained pathologist assessed the slides. A) PDXs identified as ILC. B) CTG-3434 was identified as a mixed NST ILC PDX. C) NST PDXs used for comparison.

IHC analysis for ER, PR and HER2 classified the ILC PDXs as ER+/HER2-low (N=5) and Triple-Negative Breast Cancer (TNBC) (N=2). Consistent with the HER2-low status, none of the models showed amplification of HER2 by Fluorescence In Situ Hybridization (FISH), yet they all showed HER2 staining of 1+ or 2+. The prevalence of luminal B subtypes is consistent with our recent analysis of ILC cell lines as part of the ICLE project [14], where we found the majority of cell lines to be luminal B and/or with elevated HER2 expression. It is likely that the growth *in vitro* or *in vivo* selects for cell lines and PDX that have high HER2 and growth rate. In summary, out of the 8 putative ILC PDX, 7 were confirmed as ILC and 1 was characterized as mixed NST ILC PDX (CTG-3434). The IHC analysis provided additional evidence for the lobular histology of the PDX models and their classification as ER+/HER2-low (N=5) or TNBC (N=2).

We further analyzed the WES and RNAseq data to contrast the features between the ILC (N = 7) and other NST (N = 115) PDX models (***Figure 3***). Most ILC models (5/7) were of luminal B intrinsic subtype, while the majority of NST models (68/115) were basal-like subtype (***Figure 3A, Supplementary Figure S1B***). Notably, the luminal B subtype was significantly enriched in ILC vs NST PDX models (fisher’s exact test, p = 0.005). Moreover, clustering of ILC and NST PDX models based upon the top 10% variable genes separated them into two clusters, i.e., basal and luminal/non-basal subtypes (***Figure 3B***). Most ILC models showed a similar gene expression pattern to luminal/non-basal NST PDX models. In line with the selection criteria, ILC PDX models had lower levels of *CDH1* mRNA expression vs NST PDX models (***Supplementary Figure S1C***). Similarly, differential enrichment analysis of alterations in key breast cancer genes (*TP53, CDH1, PIK3CA, ERBB2, FOXA1, MAPK31, TBX3, PTEN* and *GATA3*) between ILC and luminal/non-basal NST revealed significant (fisher’s exact test, p = 0.005) enrichment of *CDH1* alterations in ILC models, as expected (***Figure 3C***). Other top altered genes in ILC models included *TP53* (57%) and *PIK3CA* (57%) among others, while those in luminal/non-basal NST models included *TP53* (52%), *PIK3CA* (52%), *ERBB2* (42%), *GATA3* (32%) among others (***Supplementary Table S3***). Notably, ILC PDX models show a higher rate of alteration frequency in *TP53* gene, compared to clinical ILC samples (∼8% in primary tumors and ∼20% metastatic tumors) [14]. This is also true for ILC cell line models which show 88% alteration frequency in *TP53* gene [14]. The alteration frequency in *TP53* and *PIK3CA* genes was not significantly different between ILC and luminal/non-basal NST PDX models (***Figure 3C***). As expected, the top altered gene in basal NST PDX models was *TP53* (87%) (***Supplementary Table S3***) [12]. In summary, ILC PDX models were predominantly of luminal subtype, showed enrichment for alterations in the *CDH1* gene and had significantly lower *CDH1* mRNA levels than NST PDX models.

**Figure 3:**
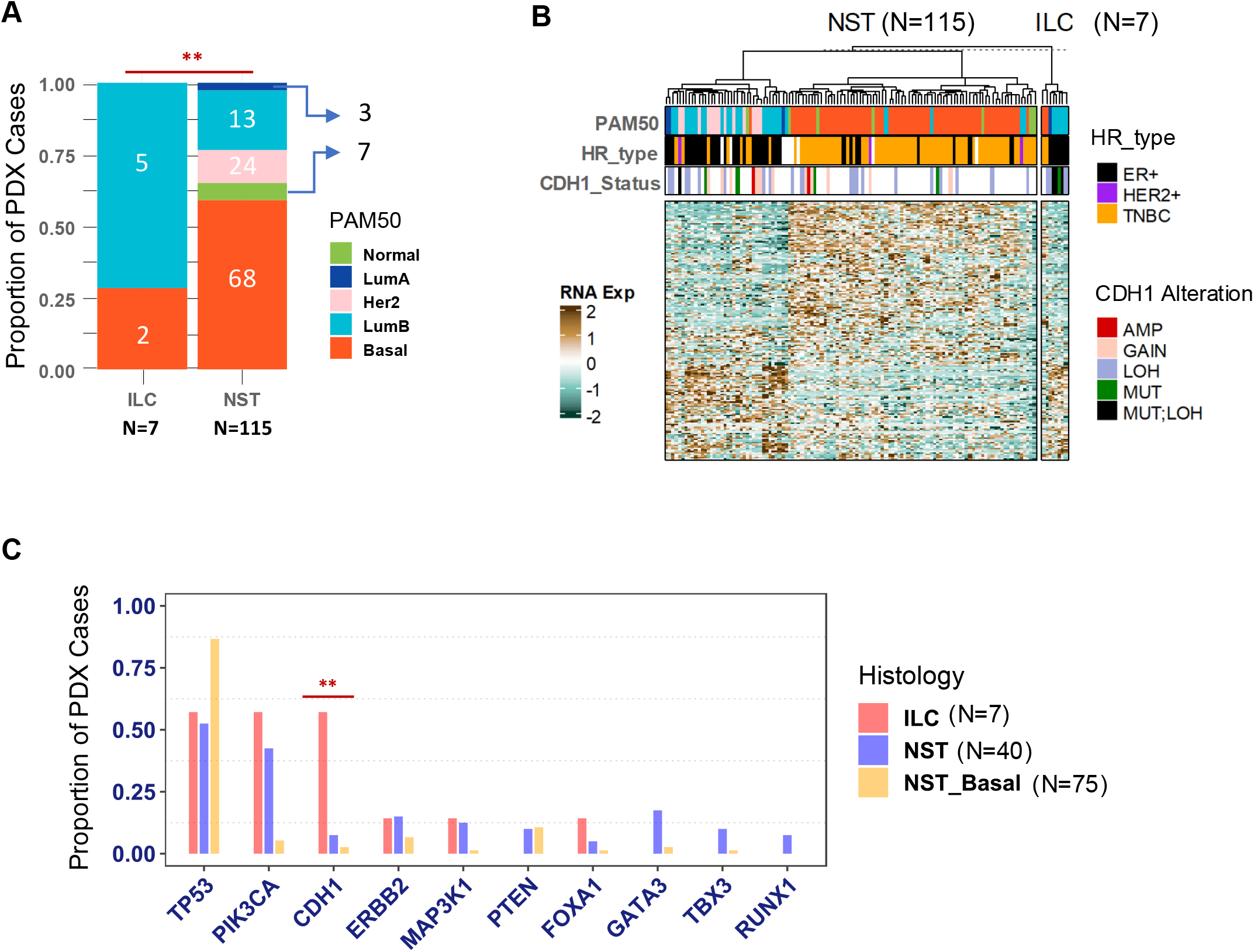
Molecular features of ILC PDX models. **A)** PAM50 subtypes in ILC vs NST/IDC PDX models. Luminal subtypes were significantly enriched (Fisher’s Exact Test, p = 0.01) in ILC vs NST/IDC PDX models. **B)** Gene expression heatmap of top 15% variable genes split by NST and ILC PDX models. **C)** Frequency of various gene alterations (MUT, MUT;LOH or MUT;GAIN) in all ILC, non-basal NST and basal NST across breast cancer genes frequently altered in breast patient tumors. *CDH1* alterations were significantly enriched in ILC vs non-basal NST (fisher’s exact test, p = 0.005) and vs. basal NST (fisher’s exact test, p = 0.0003). Other gene alterations were not significantly different between ILC and non-basal NST PDX models.

## Discussion

ILC is a unique subtype of breast cancer that accounts for 10-15% of all breast cancer but has historically been understudied, in part due to the lack of appropriate research models [15]. To address this, additional *in vitro* and *in vivo* models are needed to study ILC biology and test targeted therapies. Patient-derived xenograft (PDX) models are critical *in vivo* models for breast cancer research [15, 16]. To address this need, we utilized WES and RNAseq datasets from Champions Oncology’s Lumin portal to identify potential ILC PDX models and validated them using histopathologic analyses. The study demonstrates how existing PDX banks can be interrogated to identify models of unique histological and molecular subtypes of breast cancer.

Our study identified seven PDX models of ILC based upon truncating *CDH1* mutation and/or low CDH1 mRNA expression and confirmed using IHC and histopathological assessment. Findings from our study show that these ILC PDX models are predominantly of luminal intrinsic molecular subtype, show enrichment of *CDH1* mutations and exhibit lower levels of *CDH1* mRNA expression compared to NST, which is consistent with the characteristics of human ILC [17]. In addition to *CDH1* alterations, ILC PDX models also showed alterations in various other key breast cancer genes including *TP53, PIK3CA*, among others, while lacking alterations in genes such as *ERBB2* and *GATA3* which were more frequently altered in luminal/non-basal NST PDX models. Our findings are consistent with previous studies describing molecular characteristics of ILC [7, 17, 18].

The CTG-3434 PDX model was initially annotated as ILC and therefore included in our study. yet we identified this model as mixed NST ILC upon histopathological analysis. Following challenges in expansion of this PDX line, P5 was successfully grown and the tissue from P5 was characterized as mixed NST ILC, similar to P3 generation. The finding of mixed histology in PDX models is not unique and has been reported in other studies, for example, a study by DeRose et al. (2011) reported that some PDX models derived from breast cancer patients exhibited mixed histology, with both ductal and lobular components present in the same tumor [19]. In addition, the successful growth of the CTG-3434 PDX model in passage 5 (P5) is consistent with previous studies that have shown that PDX models can exhibit variable growth rates and success rates depending on the passage number [20]. Overall, the finding of mixed histology in the CTG-3434 PDX model, highlights the complexity of developing ILC models and underscores the importance of careful histological analysis and comprehensive characterization of PDX models.

Our study has some limitations. First, the identified ILC PDX models were based on truncating *CDH1* mutation and/or low *CDH1* mRNA expression, which may not fully represent the heterogeneity of ILC, especially those cases that do not show either *CDH1* mutations or low *CDH1* mRNA levels but lack E-cadherin expression [21]. It is possible that using additional criteria such as mutations in other adherens junction genes (e.g. *CTNNA1*) or epigenetic changes may reveal further models. Secondly, the number of identified ILC PDX models is relatively small, primarily consisting of luminal B and basal-like subtypes due to their growth advantage, rather than the more common luminal A subtype found in human ILCs. Further studies with larger sample sizes and more comprehensive characterization of ILC PDX models are needed to identify additional models for ILC and other special subtypes of breast cancer.

Overall, these findings provide additional insights into the molecular characteristics of ILC compared to NST PDX models. Understanding of molecular subtypes, gene expression features, and breast cancer gene mutations in the ILC PDX models could assist researchers in choosing the appropriate model for human ILC research and enable development of targeted therapies for this disease. Additionally, our study has implications for the development of new treatment strategies for ILC. The identification of seven ILC and one mixed NST ILCPDX models provides valuable tools for studying ILC biology and testing targeted therapies [8, 16]. Importantly, our study demonstrates how existing PDX banks with in-depth multi-omic and pathology analyses can be interrogated to identify models of unique histological and molecular subtypes of breast cancer.

## Conclusion

In conclusion, we characterized ten PDX models and identified seven ILC PDX models based upon *CDH1* mutation, low E-cadherin mRNA expression, and histopathological analysis. Our findings highlight valuable new models for studying ILC biology and testing targeted therapies in PDX models. In addition, the genomic profiling techniques used in this study provide insights into the molecular characteristics of ILC models, which can help researchers utilize these models appropriately for their research. Further studies with larger sample sizes and more comprehensive characterization of ILC PDX models are needed to validate our findings and to develop further models for ILC research.

## Acknowledgements

This study was in part supported by the National Institutes of Health (NIH) Award Number R01CA252378 and the Oncology Models Forum (to SO and AVL), P30CA047904, the Shear Family Foundation, and BCRF awards to SO and AVL. Computational resources were provided in part by the HTC cluster, supported by NIH Award Number S10OD028483.

## Author contributions

J. Hooda and J.M. Atkinson contributed equally to this work. J. Hooda: Conceptualization, supervision, investigation, visualization, methodology, writing–original draft, project administration, writing–review and editing; J.M. Atkinson: Conceptualization, supervision, investigation, visualization, methodology, writing–original draft, project administration, writing– review and editing O.S. Shah: Computational analysis, visualization, methodology, and editing; M. Yates: Investigation; D.D. Brown: Investigation; M. DeBerry: Investigation, visualization; S. Cairo: Investigation, project administration, visualization, and editing; P. Schiavini: Investigation, project administration, visualization, and editing; H.-W. Tsai: Investigation, visualization; M. Zipeto: Investigation, computational analysis; R. Bhargava: Investigation, visualization; S. Oesterreich: Resources, supervision, conceptualization, visualization, writing–review and editing; A.V. Lee: Resources, supervision, conceptualization, visualization, writing–review and editing.

## Supplementary

**Table S1:**
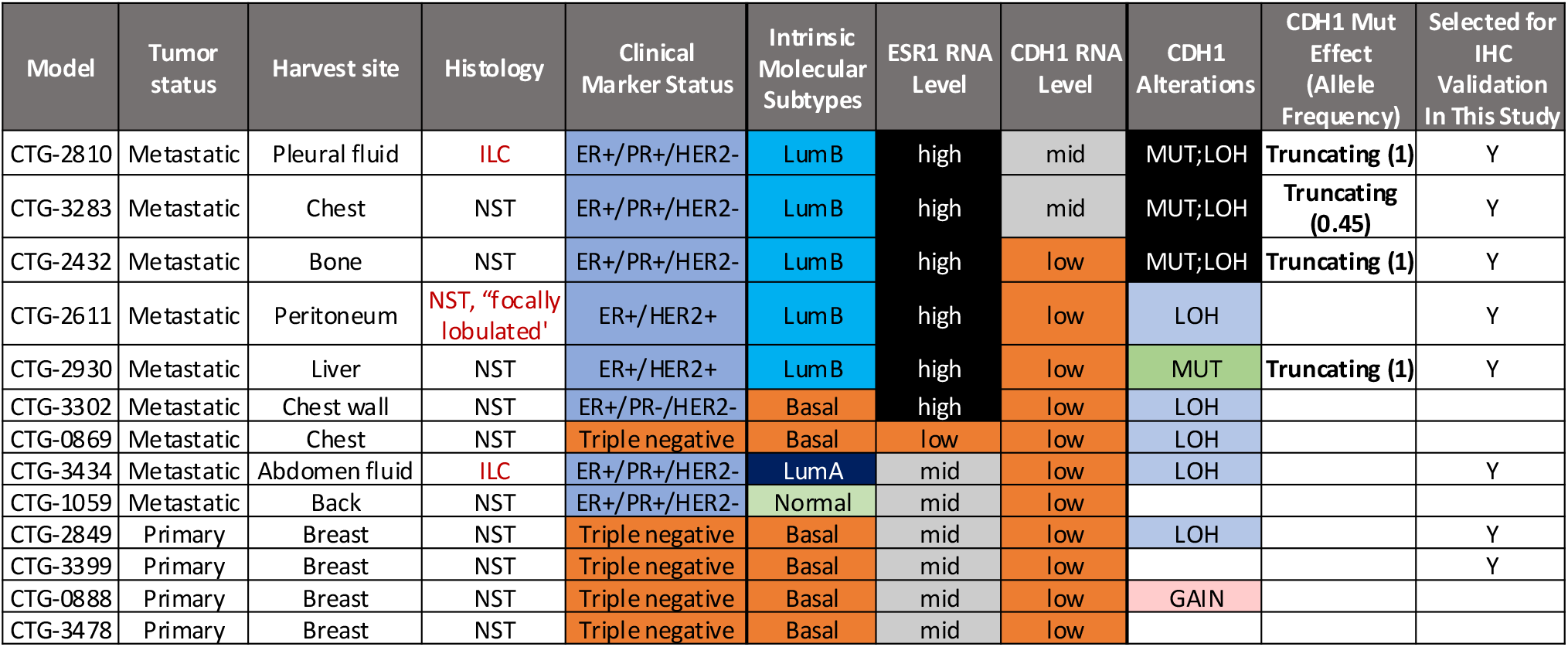
List of PDX Models (N = 13) flagged as putative ILC.

| Model | Tumor status | Harvest site | Histology | Clinical Marker Status | Intrinsic Molecular Subtypes | ESR1 RNA Level | CDH1 RNA Level | CDH1 Alterations | CDH1 Mut Effect (Allele Frequency) | Selected for IHC Validation In This Study |
| --- | --- | --- | --- | --- | --- | --- | --- | --- | --- | --- |
| CTG-2810 | Metastatic | Pleural fluid | ILC | ER+/PR+/HER2- | LumB | high | mid | MUT;LOH | Truncating (1) | Y |
| CTG-3283 | Metastatic | Chest | NST | ER+/PR+/HER2- | LumB | high | mid | MUT;LOH | Truncating (0.45) | Y |
| CTG-2432 | Metastatic | Bone | NST | ER+/PR+/HER2- | LumB | high | low | MUT;LOH | Truncating (1) | Y |
| CTG-2611 | Metastatic | Peritoneum | NST, "focally lobulated" | ER+/HER2+ | LumB | high | low | LOH |  | Y |
| CTG-2930 | Metastatic | Liver | NST | ER+/HER2+ | LumB | high | low | MUT | Truncating (1) | Y |
| CTG-3302 | Metastatic | Chest wall | NST | ER+/PR-/HER2- | Basal | high | low | LOH |  |  |
| CTG-0869 | Metastatic | Chest | NST | Triple negative | Basal | low | low | LOH |  |  |
| CTG-3434 | Metastatic | Abdomen fluid | ILC | ER+/PR+/HER2- | LumA | mid | low | LOH |  | Y |
| CTG-1059 | Metastatic | Back | NST | ER+/PR+/HER2- | Normal | mid | low |  |  |  |
| CTG-2849 | Primary | Breast | NST | Triple negative | Basal | mid | low | LOH |  | Y |
| CTG-3399 | Primary | Breast | NST | Triple negative | Basal | mid | low |  |  | Y |
| CTG-0888 | Primary | Breast | NST | Triple negative | Basal | mid | low | GAIN |  |  |
| CTG-3478 | Primary | Breast | NST | Triple negative | Basal | mid | low |  |  |  |

**Table S2:** PDX Samples Selected for IHC and Histomorphic Validation.

| PDX Model # | Collection Site | Tumor markers<br>(Champions) | Selection<br>Criteria | Reported Histology<br>(Champions) |
| --- | --- | --- | --- | --- |
| CTG-3434 | Abdominal fluid | ER+ | Low mRNA | Lobular Carcinoma |
| CTG-2432 | Bone | ER+/HER2+ | Mutation | Carcinoma |
| CTG-2611 | Peritoneum | ER+ | Low mRNA | Ductal Carcinoma<br>"focally lobulated" |
| CTG-2810 | Pleural Fluid | ER+ | Mutation | Lobular Carcinoma |
| CTG-2849 | Breast | TNBC | Low mRNA | Carcinoma |
| CTG-2930 | Liver | ER+/HER2+ | Mutation | Ductal Carcinoma |
| CTG-3283 | Chest | ER+ | Mutation | Carcinoma |
| CTG-3399 | Breast | TNBC | Mutation | Ductal Carcinoma |
| CTG-1714 | Breast | TNBC | *NST comparison* | Carcinoma |
| CTG-2518 | Breast | TNBC | *NST comparison* | Ductal Carcinoma |

**Supplementary Table S3:** Alteration Frequency of Commonly Mutated Breast Cancer Genes in basal-like NST (N=75), non-basal NST (N=40) and confirmed ILC (N=7) PDX models.

### Excel file

**Figure S1:**
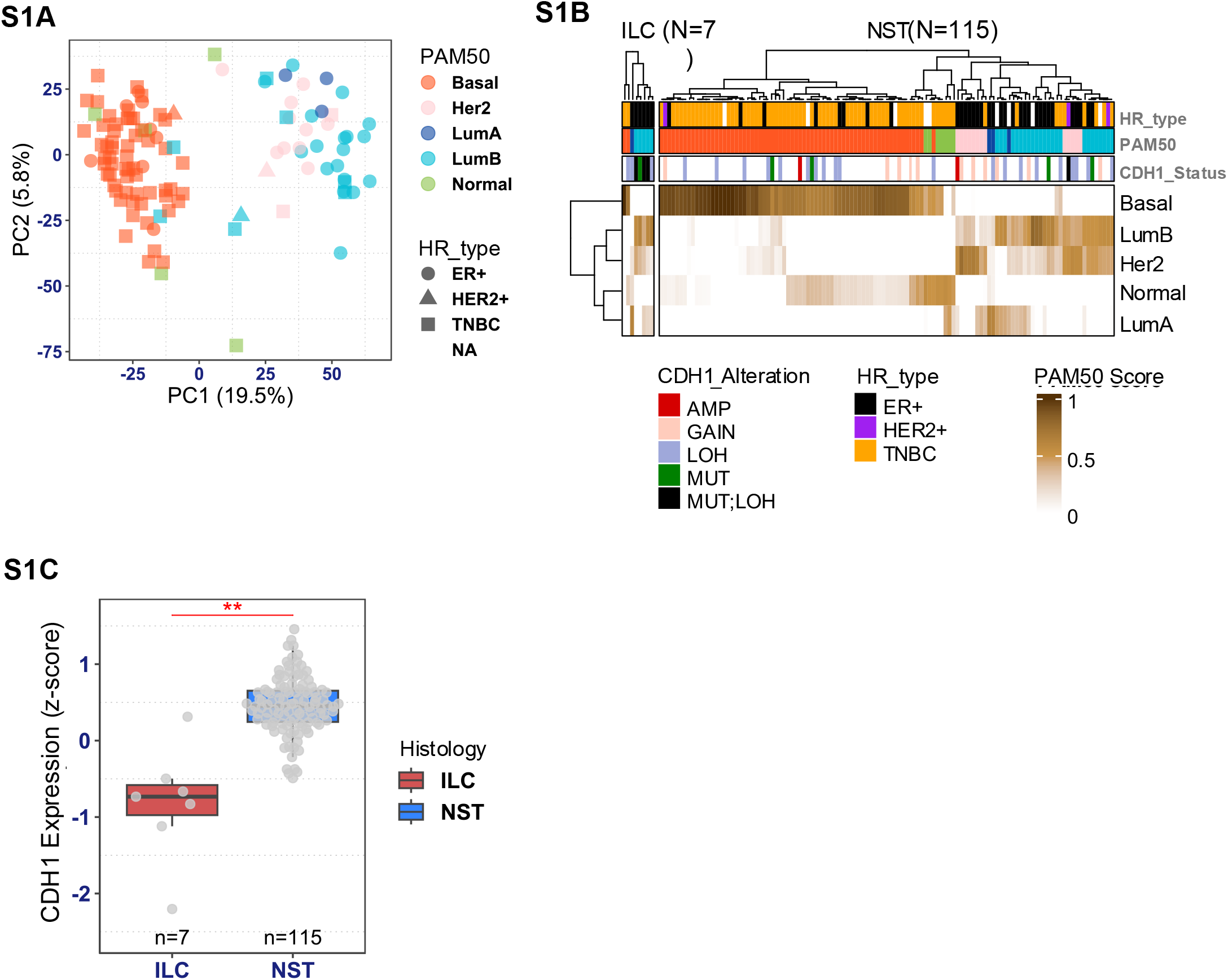
Intrinsic molecular subtypes. **A)** PCA plot of PDX models based on expression of top 10% variable genes. **B)** Intrinsic molecular subtypes in putative ILC and NST PDXs. **C)** *CDH1* mRNA expression levels in confirmed ILC PDX models vs NST. Expression difference p-value is based on t-test.

